# Sleep regulates visual selective attention in *Drosophila*

**DOI:** 10.1101/403246

**Authors:** Leonie Kirszenblat, Deniz Ertekin, Joseph Goodsell, Yanqiong Zhou, Paul J Shaw, Bruno van Swinderen

**Affiliations:** Queensland Brain Institute, The University of Queensland, Brisbane, QLD, 4072, Australia; Department of Anatomy and Neurobiology, Washington University in St. Louis, 660 South Euclid Avenue, St. Louis, MO 63110, USA

**Keywords:** attention, sleep, *Drosophila*, flies, behaviour

## Abstract

Although sleep-deprivation is known to impair attention in humans and other mammals, the underlying reasons are not well understood, and whether similar effects are present in non-mammalian species is not known. We therefore sought to investigate whether sleep is important for optimising attention in an invertebrate species, the genetic model *Drosophila melanogaster*. We developed a high-throughput paradigm to measure visual attention in freely-walking *Drosophila*, using competing foreground/background visual stimuli. We found that whereas sleep-deprived flies could respond normally to either stimulus alone, they were more distracted by background cues in a visual competition task. Other stressful manipulations such as starvation, heat exposure, and mechanical stress had no effects on visual attention in this paradigm. In contrast to sleep-deprivation, providing additional sleep using the GABA-A agonist 4,5,6,7-tetrahydroisoxazolo-[5,4-c]pyridine-3-ol (THIP) did not affect attention in wild-type flies, but significantly improved attention in the learning mutant *dunce*. Our results reveal a key function of sleep in optimising attention processes in *Drosophila*, and establish a behavioural paradigm that can be used to explore the molecular mechanisms involved.

**Summary statement:** Sleep deprivation specifically impairs visual selective attention in fruit flies, without affecting behavioural responses to simple visual stimuli.

## Introduction

The restorative effect of a night’s sleep seems obvious, yet we still know very little about the function of sleep and how it impacts our behaviour. Studies in humans and other animals suggest that a fundamental function of sleep is to preserve cognitive functions such as learning, memory and attention (Stickgold 2005). This implies that sleep promotes brain plasticity, for example by strengthening neuronal circuits to consolidate memories (Diekelmann & Born 2010), or by maintaining optimal levels of neuronal functions by globally altering synaptic strengths (Tononi & Cirelli 2013). If a major function of sleep is to promote plasticity, a brain process that may be particularly vulnerable to sleep loss is selective attention. Given that selective attention requires precise temporal coordination between different neural populations (Fries et al. 2001; Elhilali et al. 2009), it may be most vulnerable to the changes in neural processing that may accrue when sleep homeostasis mechanisms are not in place (Kirszenblat & van Swinderen 2015). Consistent with this idea, sleep-deprivation in humans leaves basic sensory processing intact (Casagrande et al. 2006; Killgore 2010; Kendall et al. 2006), whereas tasks that involve high attentional load are impaired (Chee & Chuah 2007; Kong et al. 2011). For this reason, sleep and attention may share a deeper relationship than previously thought, and one that has not been thoroughly investigated in genetic models such as *Drosophila*.

In *Drosophila*, sleep-deprivation has been associated with deficits in a variety of behaviours. Sleep-deprivation impacts olfactory and visual memory in *Drosophila* (Glou et al. 2012; Seugnet et al. 2008), as well as courtship memory (Ganguly-Fitzgerald et al. 2006) and aggression in male flies (Kayser et al. 2015). In contrast to sleep-deprivation, induction of sleep has been shown to reverse memory deficits in flies with genetic lesions and in a fly model of Alzheimer’s disease (Dissel, Melnattur, et al. 2015; Dissel, Angadi, et al. 2015). This suggests that sleep can be used as a powerful therapeutic to enhance memory. As the ability to pay attention is a prerequisite for all of the aforementioned behaviours, we questioned whether optimising selective attention may be a key function of sleep in *Drosophila*.

In this study, we developed a simple, high-throughput method to study visual selective attention in flies. We found that whereas sleep-deprivation did not affect simple visual behaviours (optomotor and fixation), sleep-deprived flies were more distractible in a visual attention task involving competing stimuli. Normal attention was restored following sleep, indicating that sleep promotes behavioural plasticity. In contrast to sleep-deprivation, we found that inducing additional sleep using the GABA-A agonist 4,5,6,7-tetrahydroisoxazolo- [5,4-c]pyridine-3-ol (THIP) had no effect on wild-type flies but could restore normal attention to *dunce* learning mutants. Together, our results suggest that sleep optimises selective attention processes in *Drosophila*.

## Methods

### Fly stocks

Strains used in this study were *Canton-S* (*CS*) wild-type flies, *dunce*^*1*^ mutants (outcrossed to CS), *rutabaga*^*2080*^, *pdf-gal4* flies (Bloomington Stock Centre, Indiana), and *UAS-Chrimson* (P[20xUAS-IVS-CsChrimson.mVenus]attp18) flies (a gift from Vivek Jayaraman, Janelia Farm Research Campus, Virginia, U.S.A).

### Sleep and visual behaviour

Our visual arena was adapted from Buridan’s paradigm (Götz 1980). Flies had their wings clipped on CO_2_, at least 2 days prior to the experiment. During the experiment, flies walked freely on a round platform, 86mm in diameter, surrounded by a water-filled moat to prevent escape. An individual fly was only tested once in each experiment, such that it was not influenced by previous visual stimuli. The temperature of the arena was 24-26 °C during experiments. Each experiment lasted 3 minutes, and the visual stimuli were presented on the horizontal or the vertical axes in alternation. Optomotor experiments were conducted with clockwise and anticlockwise gratings for 1.5 minutes each. A camera (Sony Hi Resolution Colour Video Camera CCD-IRIS SSC-374) placed above the arena was used to detect the fly’s movement on the platform at 30 frames per second, and open-source tracking software was used to record the position of the fly (Colomb et al. 2012). Sleep was quantified using the DART system as previously described (Faville et al. 2015) using the 5 minute criterion for sleep (Shaw et al. 2000; Hendricks et al. 2000).

### Visual stimuli

Each LED panel comprised 1024 individual LED units (32 rows by 32 columns) and was computer-controlled with LED Studio software (Shenzen Sinorad, Medical Electronics, Shenzen, China). The LEDs had a refresh rate of 200 Hz, ensuring there was no background flicker visible to the flies. All visual stimuli were created in Vision Egg software (Maimon et al. 2008), written in Python programming language (Ferguson et al. 2017). The walls of the arena consisted of 6 LED panels of green (520nm) and blue (468nm) LEDs that formed a hexagon surrounding the moat (29cm diameter, 16cm height), and onto which the visual stimuli were presented. Fixation stimuli were two dark stripes 180° apart, each 9° in width and 45° in height from the centre of the arena. The fixation stripes ranged from 38 - 55° in height and 4 – 14° in width depending on the fly’s position in the arena. For visual competition experiments, 7 Hz flickering stripes (targets) were overlayed on top of a 3 Hz grating (speed 54° s^-1^, luminance 402 Lux) in the background. For visual experiments in which stationary stripes were used as a distractor, the target stripes were flickered at 3 Hz, and the distractor stripes were non-flickering. For experiments in which the luminance contrast of the grating was increased, the increments used were 0, 75, 146, 224, 402, 473, 649, 730 Lux.

### Sleep-deprivation

3-5 day old female CS flies with clipped wings were sleep-deprived for 24 hours from 11am until 11am the next day, and then tested in the visual arena immediately afterwards. Mechanical sleep-deprivation was achieved using a sleep nullifying apparatus (SNAP) device, which has been shown to sleep deprive flies without triggering stress responses (Shaw et al. 2002). The device tilted back and forth, forcefully knocking and displacing flies every 20s (or 10s where indicated). Flies were contained in vials (20 females, 5 males) during the sleep-deprivation, because containing them individually in small tubes affected their performance in the attention assay. During the experiments, flies waiting to be tested were gently handled at least every 3 minutes so that they could not fall asleep prior to testing.

### Optogenetics

Flies were grown on standard (agar/yeast) medium until 2-3 days prior to an experiment, after which they were transferred to standard medium supplemented with 0.5µM all-*trans*-retinal. Chrimson was activated using Red-Orange LEDS (Luxeon Rebel, 617nm, 700mA, Phillips LXM2-PH021-0070) as previously reported (Klapoetke et al. 2014). Flies were illuminated with >1000 Lux from 4 LED arrays.

### Pharmacology

THIP was administered to flies in standard food at 0.1mg/ml concentration for 2 days, and removed 1 hour prior to measuring performance, as described previously (Dissel et al., 2015).

### Data Analyses

Analyses of visual responses were performed using CeTran (3.4) software (Colomb et al, 2012), as well as custom-made scripts in R programming language. Stripe deviation was calculated as the smallest angle between the fly’s trajectory and either of the vertical stripes (Ferguson et al. 2017; Colomb et al. 2012). For optomotor responses, the angular velocity (turning angle in ° s^-1^) in the direction of the moving grating was calculated. All sleep and arousal metrics were obtained through the DART software. Statistical analyses were performed using Prism, R and MATLAB software. Lillifors tests were performed to confirm the normality of the data, and *t* tests or one-way ANOVAs were used to detect significant differences between groups.

## Results

### A visual attention paradigm for freely walking flies

Selective attention allows us to focus on a single object or group of objects, while ignoring less salient information (La Berge 1983; Eriksen & St James 1986). To examine attention-like behaviour in freely-walking *Drosophila*, we designed a paradigm to measure behavioural responses of flies to competing visual stimuli, involving ‘targets’ and ‘distractors’. We took advantage of two robust visual behaviours in *Drosophila*: fixation – a fly’s tendency to orient and walk towards a visually salient object, and optomotor behaviour, whereby a fly will turn in the same direction as wide-field motion to stabilize its visual surroundings (Heisenberg et al. 1984). Fixation has been previously measured using Buridan’s paradigm, where flies with clipped wings walk back and forth between two opposing vertical black stripes (Götz 1980; Ferguson et al. 2017). We modified this paradigm to measure behavioural responses to competing visual stimuli, using two opposing stripes as ‘targets’ in the foreground and adding a wide-field motion stimulus as the ‘distractor’ in the background (Fig. 1A). A similar configuration has been used for ‘Fig./ground’ discrimination in tethered flight experiments (Heisenberg et al. 1984; Fox et al. 2014; Fenk et al. 2014; Aptekar et al. 2015; Aptekar & Frye 2013), but this complex stimulus has not been tested in walking flies.

**Figure 1.**
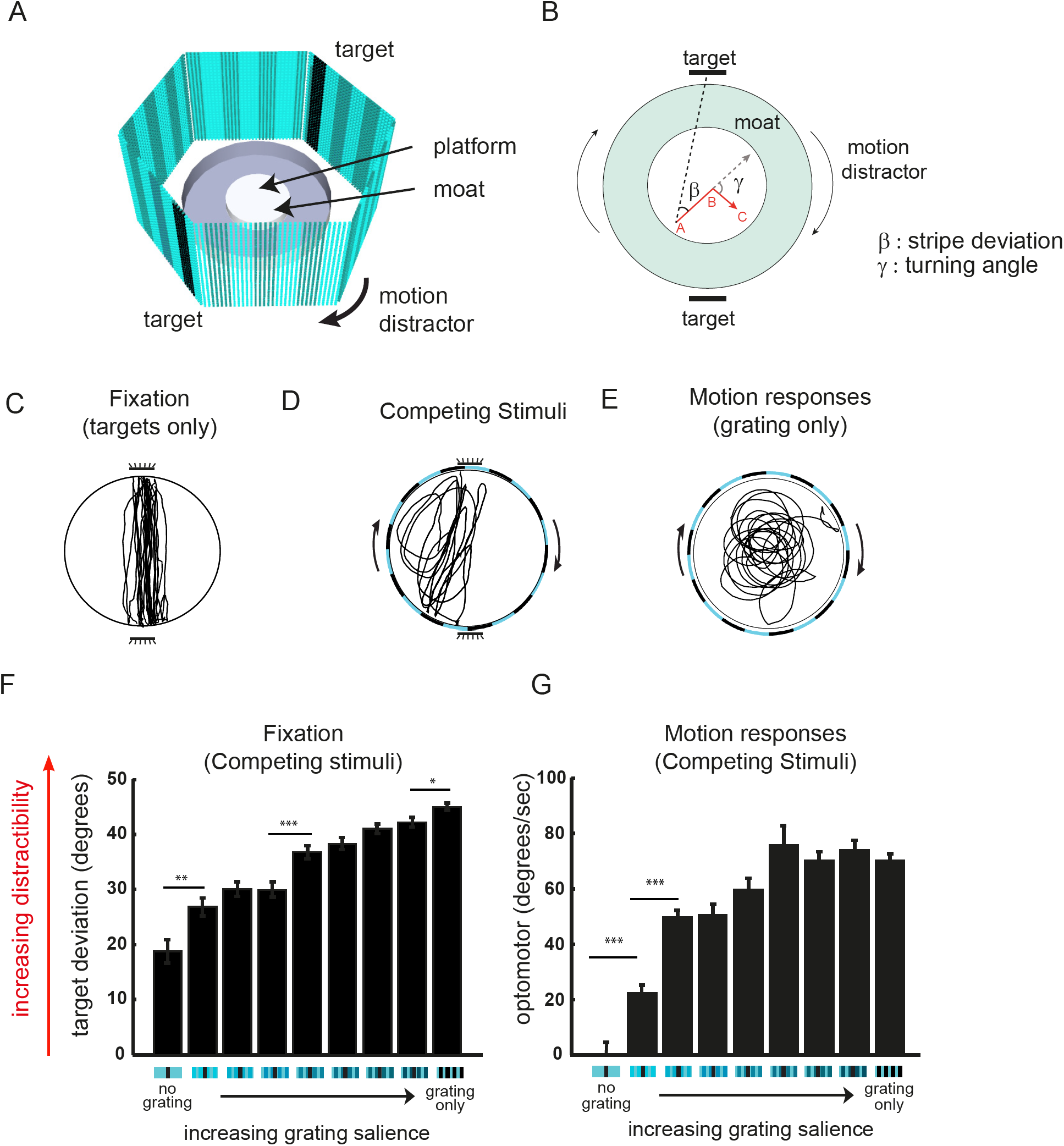
A visual attention paradigm. A) Diagram of set up. Flies walk on a platform surrounded by a moat of water, with six LED panels forming walls onto which visual cues are presented. Visual attention is measured by fixation on either of two opposing stripes (‘fixation targets’) in the presence of a motion stimulus (‘distractor’). B) Calculating fixation and optomotor responses (adapted from Colomb et al, 2013). Target fixation is measured by the stripe deviation (β), the angle between the fly’s heading direction (e.g. red line from A to B) and the centre of the target stripe that is in the direction of movement (dotted black line). Response to the motion stimulus (optomotor) is determined by the fly’s angular velocity, i.e. the fly’s turning angle (g)/second. C-E) Example traces of a fly’s walking path in response to flickering target stripes (C), a motion stimulus (D) or competing stimuli (E). F,G) Quantification of target deviation (F) or optomotor response (G) in wild type Canton-S flies when different visual cues were presented: fixation targets alone (‘no grating’) or fixation targets combined with gratings of increasing salience (luminance contrast), or grating alone. n>10 flies for each condition, *p<0.05,**p,0.01,***p<0.001 by t-tests between adjacent conditions.

We measured fixation on the targets by calculating the deviation angle between the fly’s heading direction and the target stripe, with smaller angles indicating greater attention to the target (Fig. 1B) (Colomb et al. 2012). Response to the distractor, the motion stimulus, was measured by the angular velocity in the direction of motion (the turning angle s^-1^(y): Fig. 1B). First, we investigated visual responses to both stimuli alone, across a range of frequencies (Fig. S1A,B). Flies fixated best on targets flickering in the range of 3 – 7 Hz (Fig. S1A), while optomotor responses were highest at 16 Hz (Supplementary Fig. 1B). For visual competition experiments, we used a target frequency of 7 Hz, as this stimulus was very salient to the flies and has previously been found to evoke strong fixation behaviour (Paulk et al. 2015). The distractor was a lower contrast grating that spanned the entire arena and moved “behind” the flickering targets. We used a grating frequency of 3 Hz (54 ° s^-1^) at a mid-luminance contrast, since we found responses under these conditions were robust and consistent among flies, eliciting circular walking responses (Fig. 1E), and provided a low enough level of distraction for the flies such that they could still continue to fixate on the targets. Adding this low-contrast motion distractor significantly reduced flies’ fixation on the target stripes: the straight paths evident when the targets were presented alone (Fig. 1C) were replaced by a combination of straight paths and circular paths, often angled in the direction of motion (Fig. 1D). We modified the salience of the grating distractor by adjusting the luminance contrast in linear increments (see methods) (Fig. 1F,G). As expected, when the grating had a higher contrast (more salient), the distraction effect increased (Fig. 1F). Motion responses to the distractor also increased (Fig. 1G) but appeared to plateau earlier than target deviation in response to increasing the distractor salience. This suggests that increased target deviation (Fig. 1F) may be a more sensitive measure of the distractor effect than increased optomotor behaviour (Fig. 1G). In summary, our results show that attention-like responses towards fixed targets can be effectively modulated (and titrated) by a motion distractor stimulus in freely walking flies.

### Sleep-deprived flies are more distractible

Studies in humans and other animals have shown that sleep loss can impair a variety of cognitive behaviours, such as learning, memory and attention (Alhola & Polo-kantola 2007; Drummond et al. 2012; Lim & Dinges 2010). Similarly, studies in *Drosophila* have indicated a role for sleep in visual and olfactory memory (Glou et al. 2012; Li et al. 2009; Seugnet et al. 2008; Seugnet et al. 2011). However, it is not known whether sleep modulates attention-like behaviour in *Drosophila*. To investigate whether sleep-deprivation affects visual attention, we sleep-deprived flies for 24 hours using the Sleep Nullifying Apparatus (SNAP) according to previous methods (Shaw et al. 2002). We first confirmed that flies were sleep-deprived by examining their sleep rebound (*i.e.* an increase in sleep need following sleep-deprivation). Following sleep-deprivation, sleep was monitored using the DART system (Faville et al. 2015) across three consecutive days, while another group of flies had their attention tested (Fig. 2A). As expected, sleep-deprived flies slept more than control flies the day after sleep-deprivation (day 1) particularly during the daytime, whereas by days 2 and 3 there was no difference in sleep duration between the sleep-deprived and control flies (Fig. 2B).

**Figure 2.**
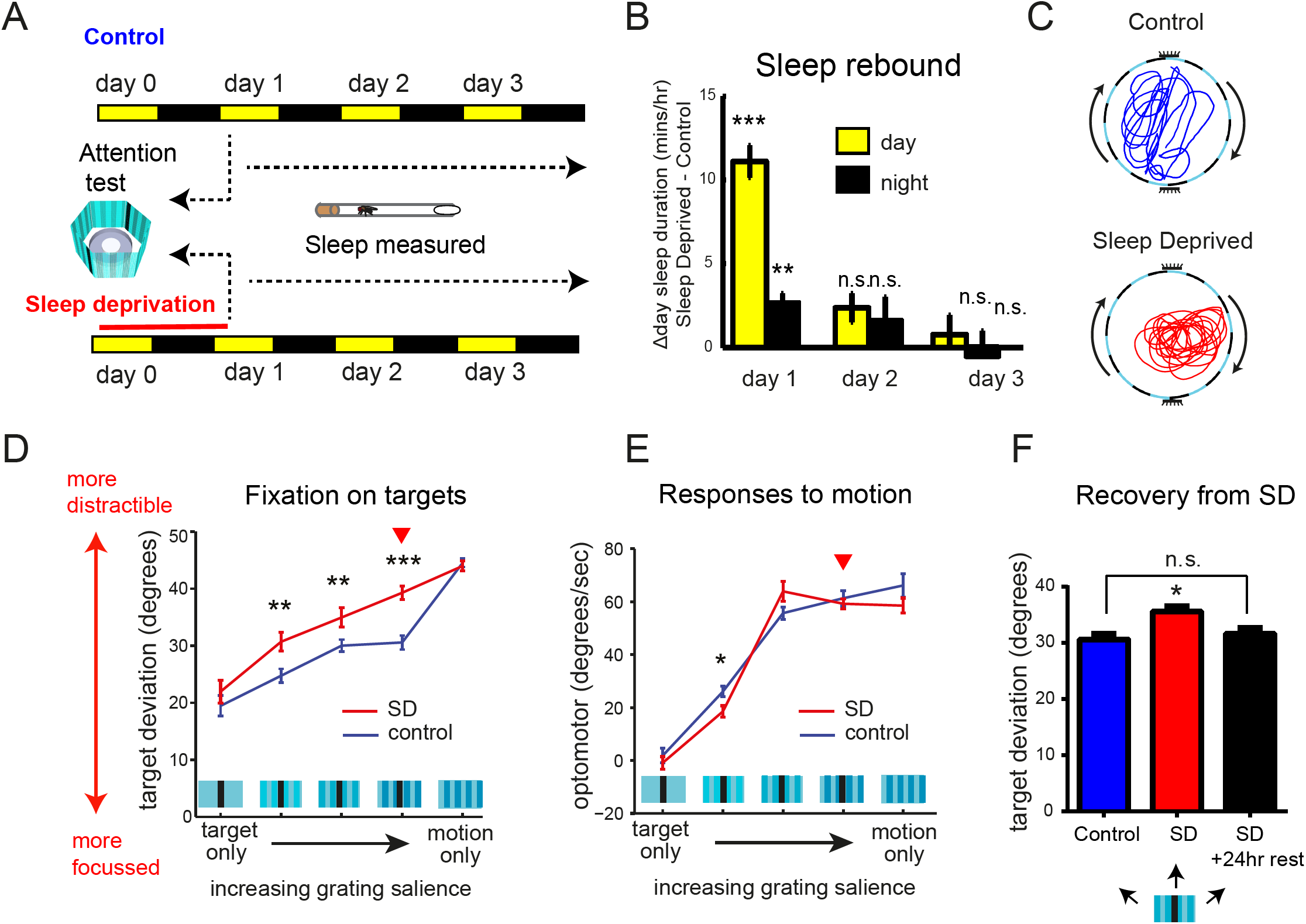
Sleep deprivation makes flies more distractible. A) Sleep deprivation protocol. Flies were sleep deprived for 24 hours and visual attention was tested the following day (day 1), and compared to control (non-sleep deprived flies). B) Flies subjected to the sleep deprivation protocol in (A) showed a sleep rebound (increase in sleep) measured by the difference in sleep compared to control (non-sleep deprived) flies. C) Example traces of walking behaviour in a control and sleep deprived fly responding to competing visual cues. D,E) Quantification of target deviation (D) or optomotor response (E) in control vs sleep deprived flies responding to target objects alone (target only), competing visual cues (objects and grating together, with three levels of grating salience), and grating alone (motion only). F) Attention returns to normal levels in flies rested for 24hrs following sleep deprivation (tested with objects and mid-salience grating). n= 60 flies in (A,B) n>16 flies for each condition in (D,E) and n>12 flies in (F). Red arrowheads indicate the grating contrast that was used for all subsequent experiments *p<0.05 (one-way Anova in F, t-test in E) and **p <0.01, ***p <0.001 by t test between control and sleep deprived. Error bars show the s.e.m.

Following sleep-deprivation we measured fixation to stationary targets and optomotor responses to a moving grating, in scenarios in which the stimuli were presented alone or in competition (with the grating at three different levels of luminance contrast of the distractor) (Fig. 2D,E). Optomotor responses to the moving grating in all visual scenarios were largely unaffected by sleep-deprivation (Fig. 2E), as were fixation responses in the absence of the grating (Fig. 2D, first point from the left). Interestingly, the effect of sleep-deprivation on fixation behaviour was only evident when the two visual stimuli were combined (Fig. 2C), as indicated by significantly increased target deviation for all three combined conditions (Fig. 2D). This suggests that sleep-deprived flies are specifically affected in their response to visual competition. Finally, we tested whether flies could recover from sleep-deprivation by allowing them to rest for 24 hours. For this experiment (and all subsequent experiments involving visual competition), we used a grating of mid-luminance contrast (indicated by the red arrow heads in Fig. 2E,F). Again, we found that sleep-deprived flies were more distractible, but their attention returned to normal following 24 hours rest (Fig. 2F). Overall, our results suggest that sleep-deprivation alters visual attention, and that this effect is reversible.

Although we saw no alterations to optomotor responses of sleep-deprived flies when the grating was presented alone, we considered the possibility that the sleep-deprivation phenotype may be caused by an increased sensitivity to motion, which could potentially arise from the preceding 24hours of constant motion they experience in the SNAP apparatus. However, flies sleep-deprived in 24hours of darkness showed a similar impairment of attention compared to controls, suggesting that visual experience did not play a role in the sleep-deprivation phenotype (Fig. S2). We next asked whether the increased distractibility of sleep-deprived flies was specific to the motion stimulus, or whether it could apply to other types of visual distractors. We used a stationary stripe as a distractor that was identical to the target but less salient as it was non-flickering (Fig. 3A). As with the moving grating in the preceding experiments, attention to the target could be titrated by modulating the salience (*i.e.,* contrast) of the distractor (Fig. 3B,C). Interestingly, increasing the salience of the distractor correspondingly increased distractibility (Fig. 3C).Furthermore, consistent with our previous attention experiments, we found that sleep-deprived flies similarly paid less attention to the target in the presence of this alternative distractor object (Fig. 3D). Our observation that sleep-deprivation is similarly disrupted by different types of distractors suggests that a common attention mechanism is affected, which is not dependent on lower level visual responses.

**Figure 3.**
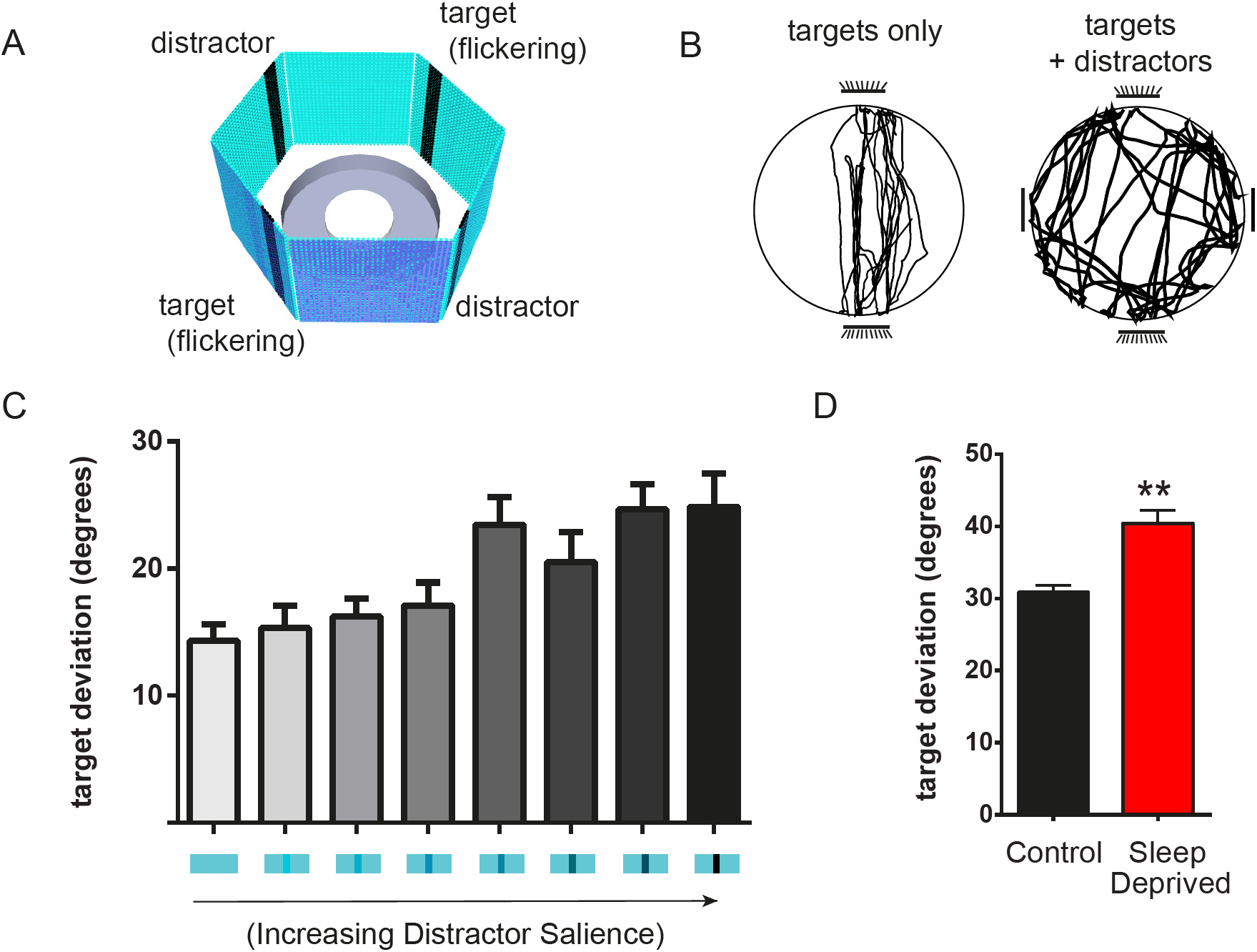
Sleep deprivation increases distractibility to stationary objects. A) Diagram of visual stimuli, with flickering target stripes and non-flickering distractor stripes. B) Example traces of a flies responding to targets only, or targets with distractor stripes. C) Target deviation scores with increasing distractor salience (luminance contrast to background). D) Target deviation for controls and sleep deprived flies in the presence of distractor objects. n<20 flies in (C) and 50 flies in (D). **p<0.001 by t-tests between adjacent groups in (C) and sleep deprived and controls in (D).

Given that sleep-deprivation reduces overall activity, we wondered whether this may explain the reduced orientation to salient features in the visual attention assay. To investigate this, we performed a correlation analysis on all data, to see if walking speed of individual flies may predict selective attention behaviour (target deviation). However, we found no correlation between attention and walking speed for either sleep-deprived or control groups (Fig. 4A,B).

**Figure 4.**
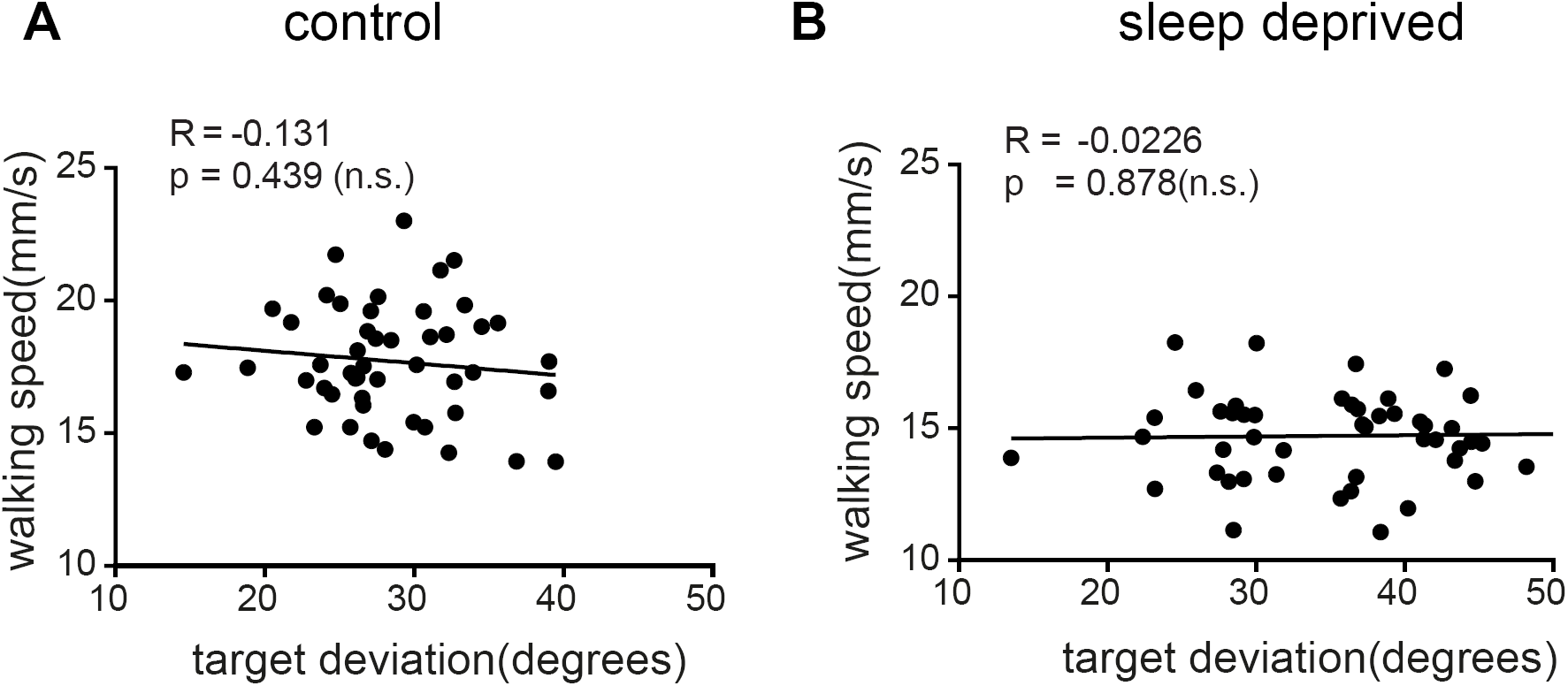
Target deviation in the attention paradigm is not correlated with walking speed or sleep following sleep deprivation. A,B) Correlation of walking speed and target deviation in control (A) and sleep deprived (B) flies. n = 48 flies. Pearson’s correlation coefficient (‘R’) were not significant (shown on graphs, alongside p values), indicating no correlations were found.

Considering that the sleep-deprivation protocol might be stressful (it involves waking the flies every 20 seconds by mechanical stimulation), we further examined the effects of different types of stress on the flies’ visual attention. This allowed us to answer two questions: 1) whether attention phenotypes could be modulated by environmental stressors (e.g. heat, mechanical stress and starvation), which would indicate that the sleep-deprivation effect on attention may simply be caused by stressful situations, and 2) whether the sleep-deprivation phenotype occurs under different environmental conditions *(i.e.,* how robust the effect is). For all these experiments we measured flies’ visual responses in the combined stimulus paradigm (in which flies fixate on targets in the presence of distractors) (Fig. 1A).

First, we measured the effect of heating flies to 31°C for 24 hours (Fig. 5A). Prolonged exposure to heat had no effect on attention, whereas the increase in distractibility caused by sleep-deprivation was still present under conditions of heat stress (Fig. 5A). This indicates that heat does not influence attention in this paradigm, or the sleep-deprived effects on attention. Next, we measured the effects of mechanical stress using a stimulation protocol that would provide the same amount of stimulation experienced during the sleep-deprivation protocol, but including rest periods for sleep. This was achieved by subjecting flies to mechanical stimulation for one hour followed by rest for one hour, repeated across 24 hours, but with double the rate of stimulation (to keep the actual number of stimuli the same as for the sleep-deprivation regime) (Fig. 5B). Unlike sleep-deprived flies, flies subjected to this stimulation regime containing rest periods had normal attention, indicating that the sleep-deprivation effect was not due to mechanical stress of perturbing the flies (Fig. 5B). We next assessed the effects of starvation, as sleep is known to be affected by food availability (Siegel et al, 2009). In *Drosophila*, food deprivation has been found to suppress sleep (Keene et al. 2010). Feeding was assessed during the sleep-deprivation by measuring the intake of food containing blue food dye and, as previously reported, sleep-deprived flies showed similar food intake to controls (Thimgan et al. 2010). We then tested whether depriving flies of food, with or without sleep-deprivation, affected visual attention. The visual attention scores of starved controls were not different from those of fed controls, suggesting that starvation *per se* does not impair visual attention (Fig. 5C). As expected, sleep-deprivation impaired visual attention in both starved and fed conditions but, interestingly, starvation was able to partially suppress this effect (Fig. 5C, “SD fed” vs “SD starved”), suggesting that starvation may protect against the detrimental effects of sleep-deprivation. In summary, our results show that sleep-deprivation consistently affects visual attention, even under conditions of heat and starvation, but that visual attention is resistant to the effects of a variety of stresses in the absence of sleep-deprivation. Therefore, sleep is more important than lack of stress for maintaining optimal levels of visual attention.

**Figure 5.**
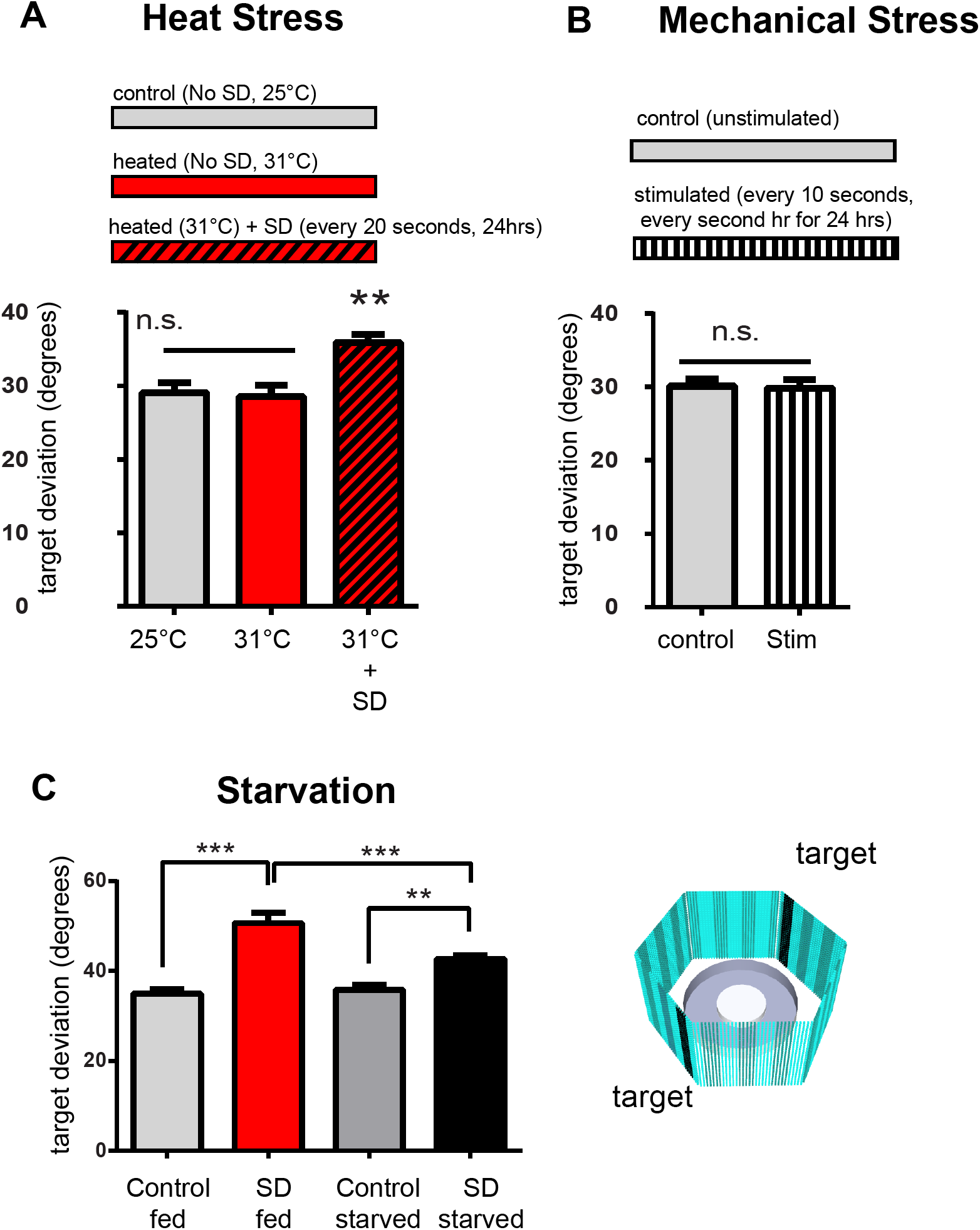
Starvation, heat, and mechanical stress effects on visual attention. Visual attention was measured by orientation towards 7hz flickering object (’targets’) with a mid-luminance contrast grating distractor (see methods). A) Target deviation in flies that underwent heat stress (31 degrees) for 24 hours with or without sleep deprivation. B) Target deviation following 24hrs mechanical stimulation. C) Target deviation in flies that were starved and/or sleep deprived for 24hrs. “Control fed” vs “control starved” groups were not significantly different. n>17 flies in (A), n>28 flies in (B) n = 20 flies in (C). **p<0.01, ***p<0.001 by One-way ANOVA. Error bars indicate the s.e.m.

Having found that sleep-deprivation affects visual attention, we also sought alternative methods to sleep deprive flies, to confirm that the sleep-deprivation effect on attention was not specific to our mechanical stimulation protocol. Pigment Dispersing Factor (PDF) neurons in the optic lobes of the fly brain are known to promote wake (Parisky et al. 2008). We hypothesised that activating these neurons for 24 hours would have wake-promoting effects that may also impair visual attention. To test this, we transiently expressed a red-light activated channelrhodopsin, Chrimson (Klapoetke et al. 2014), in the Pigment Dispersing Factor (PDF) neurons, such that exposing flies to red light could trigger wakefulness. We noticed that red light exposure caused a general decrease in sleep among all flies (Fig. 6A). However, sleep loss was greater in flies that expressed the channelrhodopsin (*pdf-gal4/+>uas-chrimson/+*) and were exposed to red light for 24 hours, as they lost more than 55% of their sleep compared to non-exposed controls. When we subsequently tested these flies for visual attention, we found that this method of sleep-deprivation resulted in an increase in target deviation specifically in the flies that were kept awake genetically, *pdf-gal4/+>uas-chrimson/+* (Fig. 6B,C). This provides further support for the idea that sleep-deprivation increases distractibility, and is not specific to the method of sleep-deprivation.

**Figure 6.**
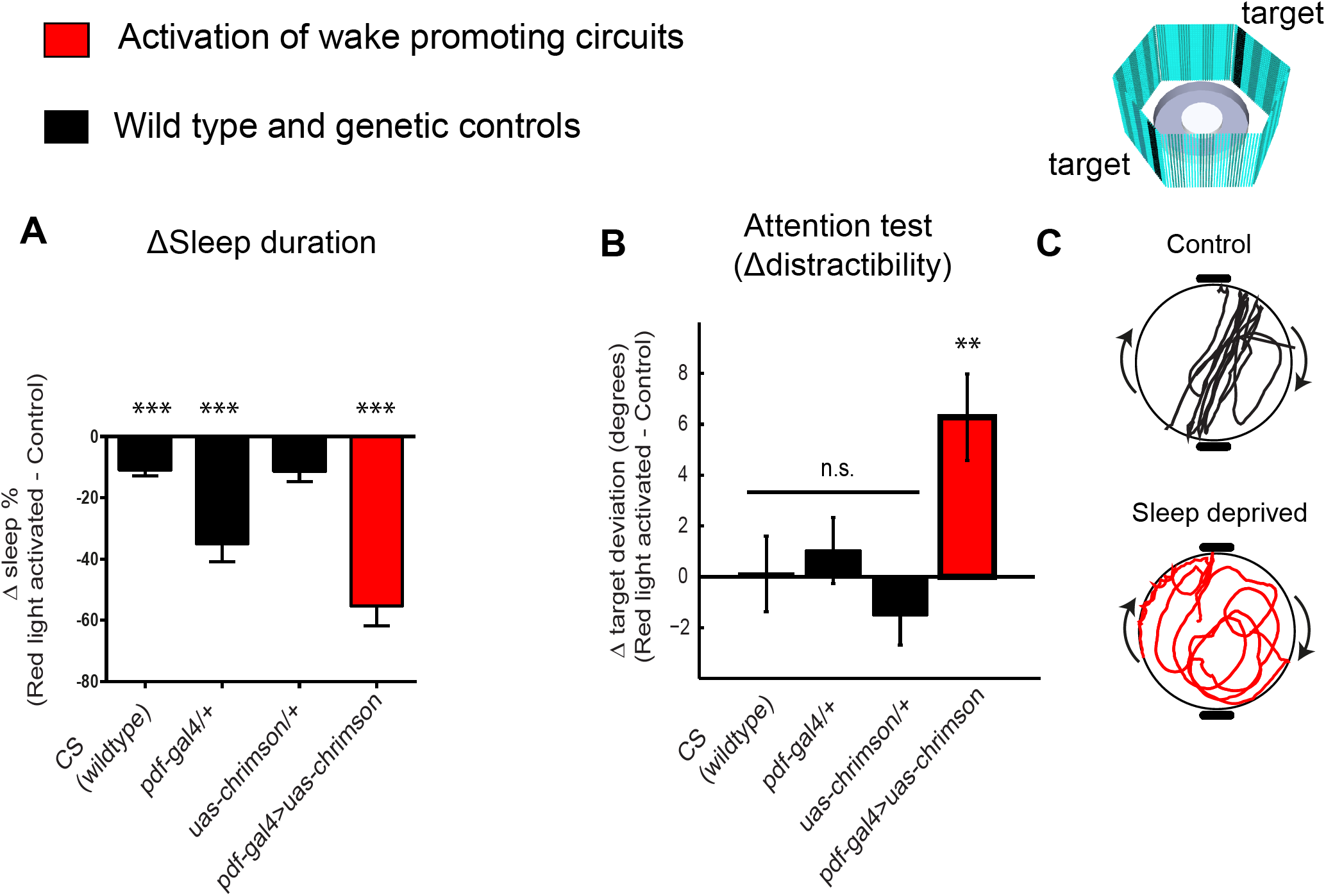
Activation of wake promoting circuits for 24 hours increases distractibility. A) Change in sleep caused by red-light exposure. n= 17 B) Change in attention (target deviation) following 24 hrs red light exposure. n = 10. C) Representative traces of a control fly (CS) compared to a sleep deprived fly (pdf-gal4>uas-chrimson) following red light exposure. t tests compared red-light activated to non-activated controls. **p=0.002, ***p<0.001, one-way ANOVA. Error bars show the s.e.m.

### Additional sleep improves attention in *dunce* mutants, but not wild-type flies

Considering that sleep-deprivation made flies more distractible, we next asked whether *increasing* sleep had the opposite effect on visual attention, making flies *less* distractible. The GABA-A agonist (THIP) has recently been used to induce sleep in *Drosophila* (Dissel, Angadi, et al. 2015; Yap et al. 2017). This increase in sleep was shown to reverse memory deficits in *Drosophila* learning mutants *dunce* and *rutabaga* (Dissel, Angadi, et al. 2015) and in a *Drosophila* model of Alzheimer’s disease (Dissel et al. 2017). As *dunce* and *rutabaga* mutants have previously been found to have defective attention processes (van Swinderen & Brembs 2010; van Swinderen 2007), we wondered whether their attention deficits could also be rescued by induced sleep.

We first quantified sleep in *dunce* and *rutabaga* mutants, and found that both these mutants slept significantly less than wild-type flies, with *dunce* flies sleeping less during the day and the night, and *rutabaga* mutants sleeping less during the day (Fig. 7A,B). THIP could then be effectively used to increase sleep in both wild-type and mutant flies, to a similar level (Fig. 7C,D). We next tested whether inducing sleep affected the visual attention of wild-type flies in our free-walking paradigm. THIP was fed to wild-type (CS) flies for two days and removed one hour prior to testing their behaviour (the same procedure used by (Dissel, Melnattur, et al. 2015). We performed a within-group experiment (the same flies tested before and after induced sleep), and a between-group experiment (aged-matched flies, with or without induced sleep). In wild-type flies we observed that distractibility (deviation away from the targets) decreased slightly for those flies that that had been induced to sleep more, in both experiments; however, this effect was not significant (Fig. 7E, p=0.084 and p=0.187 for within group and between group comparisons, by t-test.).

We next tested whether increasing sleep altered attention in *dunce* and *rutabaga* mutants. Under normal conditions (without increasing sleep), *dunce* and *rutabaga* flies were significantly less attentive towards the visual targets compared to wild-type flies as measured by increased target deviation (Fig. 7F, black bar (WT) compared to light blue bar (*dnc* (-)) and yellow bar (*rut* (-))). Interestingly, THIP administration was able to significantly reduce target deviation in *dunce* mutants but not *rutabaga* mutants (Fig. 7F), such that attention in *dunce* mutants with induced sleep was not different from that of wild-type flies (Fig. 7F, dark blue bar (*dnc* (+)) compared to black bar (WT(-)). This suggests that increasing sleep may improve attention in some learning mutants but not others.

**Figure 7.**
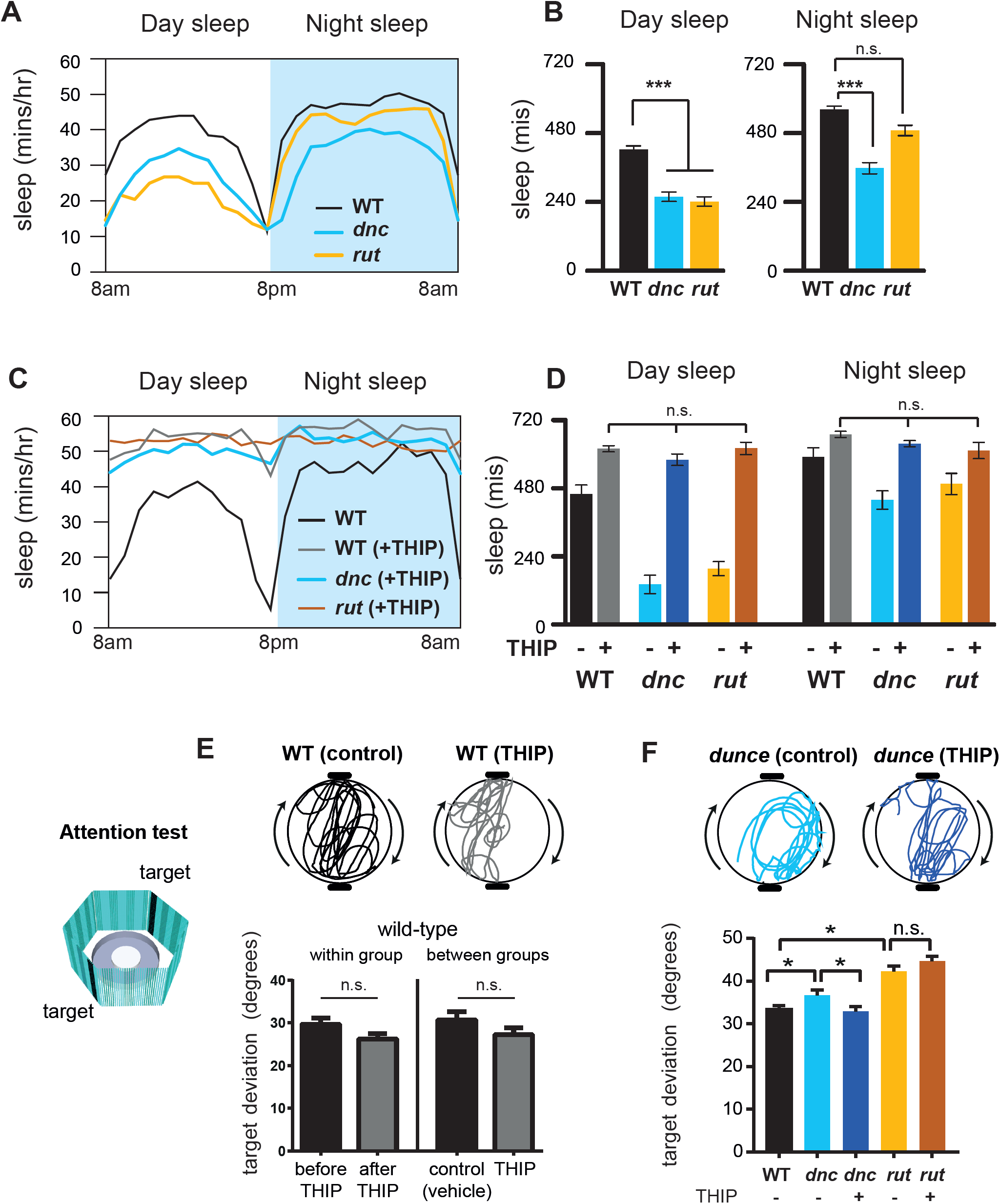
Increasing sleep improved attention in *dunce* mutants. A) Hourly sleep profiles averaged across 24 hours for wild-type (Canton-S), dunce and rutabaga flies (n>50 flies). B) Quantification of sleep duration averaged across the day and the night, from data in (A). C) Sleep across 24 hours in wild-type flies, dunce and rutabaga mutants treated with the sleep promoting agent, THIP. Wild-type sleep without THIP is shown for comparison (black trace). n>20 flies. D) Quantification of day-time and night-time sleep in wild-type, dunce and rutabaga flies with and without THIP. n>20 flies. E) Target deviation of wild-type (Canton-S) flies on control (vehicle) or THIP (0.1mg/ml) food. THIP was administered for 2 days and flies were removed from THIP 1 hr prior to the visual attention test. ‘Within group’ compared flies before and after THIP administration (n=11 flies), and ‘between group’ compared age-matched flies fed control (vehicle) or THIP food (n = 8 flies). F) Stripe deviation in dunce mutants before and after 2 days of THIP (n>30 for dunce and CS, n> 16 for rutabaga). *p<0.05, ***p<0.0001 by one-way ANOVA (A-D) and t-test (E-F). Error bars show the s.e.m.

## Discussion

Selective attention is crucial for discriminating important from irrelevant stimuli in our environment, even for insects such as *Drosophila*. The selection and suppression dynamics required for attention seem to have emerged in brains at the same time at which sleep became important for maintaining attention-related behaviours (Kirszenblat & van Swinderen 2015). In other words, some sleep functions may have evolved to optimise attention. Here, we have developed a free-walking attention paradigm which allowed us to obtain a functional readout of the effects of sleep-deprivation and sleep induction on visual attention in *Drosophila*.

Selective attention must require a certain level of brain coordination to deal with competing stimuli. This is because different stimuli may be processed by different brain regions; in our attention paradigm, the optomotor and fixation responses that compete with each other have been found to be driven to some degree by independent visual circuits (Bahl et al. 2013; Fenk et al. 2014). Furthermore, attention-like behaviour in the fly is associated with increased coherence between brain regions (Paulk et al. 2015), suggesting greater synaptic coordination. If sleep is required to maintain synaptic coordination across the brain, it follows that tasks involving greater cognitive load may be more vulnerable to sleep loss. Indeed, we found that although sleep-deprived flies showed normal responses to simple visual stimuli, they had altered responses to visual competition.

One might still ask, whether sleep has a privileged role in regulating selective attention relative to other high order cognitive functions. Sleep loss is known to affect a variety of complex cognitive behaviours in humans, such as learning and memory, creative thinking, and even the ability to speak clearly or to appreciate humour (Harrison & Horne 1997; Killgore 2010; Kendall et al. 2006). Attention is probably integral to these complex processes because it allows us to filter out irrelevant information and to select the right actions. In light of this, we would suggest that selective attention is a key *mechanism* that is affected by sleep loss, which disrupts the brain’s ability to prioritize competing information.

The sleep-deprivation effects we observed on attention seemed to be remarkably robust under different environmental conditions, as flies were still more distractible following sleep-deprivation in conditions of heat stress, darkness and starvation. Interestingly, although starvation and heat alone can promote waking in flies (Ishimoto et al. 2012; Keene et al. 2010), they did not appear to affect our attention readout, unlike the mechanical perturbation method of sleep-deprivation which is in standard use for *Drosophila*. It is possible that because starvation and heat are environmental stimuli that a fly encounters in natural situations, it has already evolved physiological protective mechanisms to cope with these kinds of stresses. This is supported by reports that heat stress response factors can protect against sleep-deprivation-induced lethality (Shaw et al., 2002), and that starvation disrupts sleep homeostasis (Keene et al. 2010; Thimgan et al. 2010). *Foraging* mutants of *Drosophila*, which tend to explore further for food, also appear to have reduced sleep need and are resistant to memory impairments caused by sleep loss (Donlea et al. 2012). More recently, it was also discovered that sexual arousal in male flies can suppress the need for sleep (Beckwith et al. 2017), and vice versa, sleep can inhibit male sexual behaviour (Chen et al. 2017). Overall, these studies suggest that sleep need is flexible and may compete with other survival needs such as food, sexual reproduction and the need to escape unfavourable environments (e.g. high temperatures). In future it would be interesting to further investigate how a fly’s environment influences its attention.

Interestingly, *increasing* sleep was able to improve the attention of *dunce* mutants, but not wild-type flies or *rutabag*a mutants. The finding that wild-type flies did not show improved attention following induced sleep is consistent with previous reports that increasing sleep does not improve learning and memory in wild-type flies (Dissel, Angadi, et al. 2015). One interpretation is that attention is already optimal in wild-type *Drosophila*. In contrast, both *rutabaga* and *dunce* mutants have previously been identified as having attention deficits (van Swinderen 2007), and our result confirmed this finding in a free-walking attention paradigm. However, it is not clear why inducing sleep would improve attention specifically in *dunce* mutants, and not in *rutabaga* mutants. It is possible that this may relate to their different sleep phenotypes – *dunce* mutants appear more severely sleep-deficient in our DART system (including at night, which *rutabaga* mutants are not) meaning that THIP had a greater ability to restore sleep to *dunce* mutants. Related to this, it is possible that the attention deficits of these mutants are due to different underlying causes. For example, the mutants may have poor attention because they sleep less, or they may sleep less because they have poor attention.

How inducing sleep could improve attention, or other aspects of cognition such as learning and memory, also remains unclear. There is some evidence that the sleep-promoting agent THIP can interact with specific GABA-receptors to promote sleep in flies, via known sleep-promoting neurons of the dorsal fan-shaped body (Dissel, Angadi, et al. 2015), but can also modulate dopaminergic pathways to facilitate memories (Berry et al. 2015). Interestingly, a recent study found that increasing sleep promoted survival of wild-type flies exposed to oxidative stress (Hill et al. 2018). Whether increasing sleep in flies improves brain function through specific circuits, or by general cellular mechanisms like reducing reactive oxygen species (ROS) needs to be further investigated. Our study opens up an opportunity to understand the molecules, circuits and environmental factors involved in the role of sleep in optimising attention.

## Acknowledgements

We would like to thank Rowan Tweedale for comments on the manuscript. We would like to thank John John for help with behavioural experiments, and Richard Faville and Ben Kottler for help with DART software. This work was supported by an NIH grant RO1 NS076980-01 to PJS and BVS and an Australian Research Council grant DP140103184 to BVS.

---

**Figure S1.**
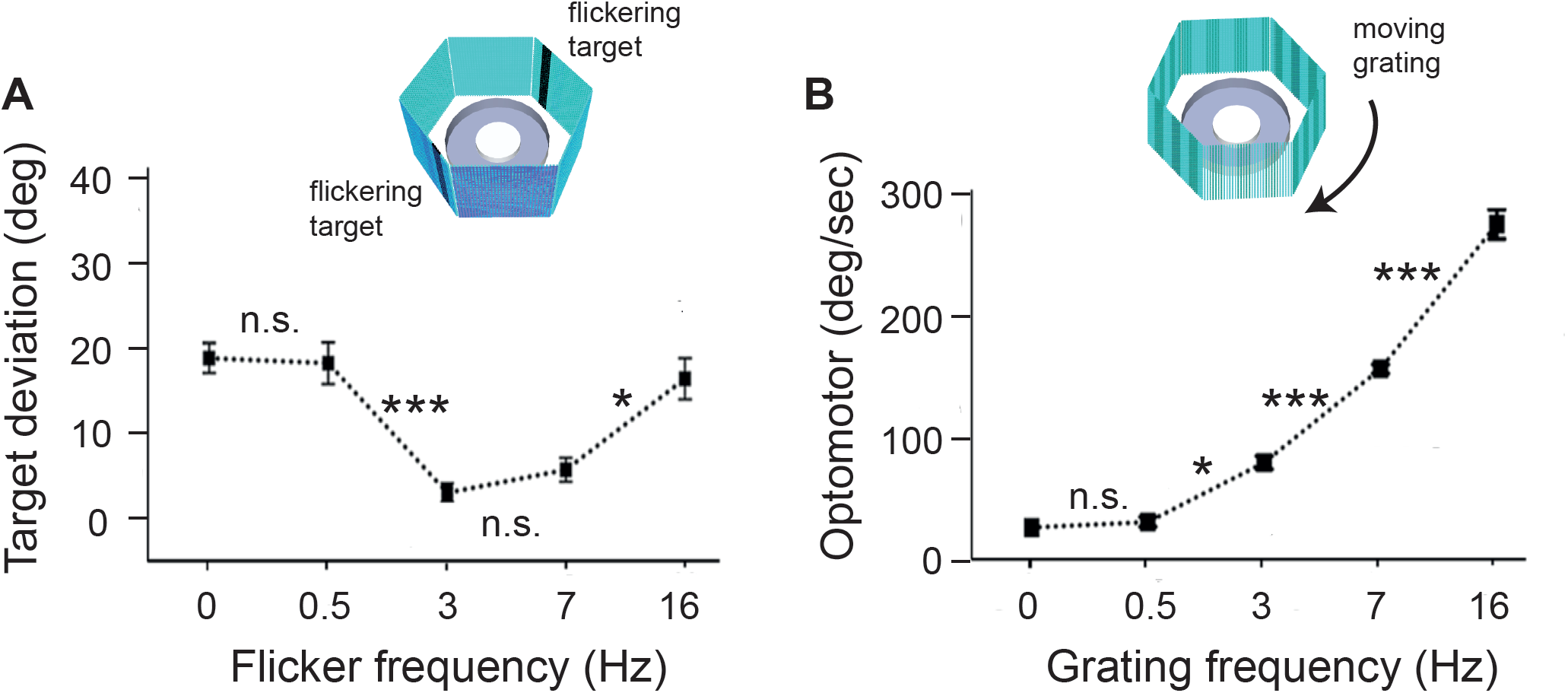
Visual responses are modulated by flicker and grating frequency. (A) Visual fixation in wild-type flies responding to fixation targets that were static (0 Hz) or flickering at 0.5 Hz, 3 Hz, 7 Hz, or 16 Hz. Lower target deviation indicates greater fixation. n = 14 flies for each condition. Asterisks indicate significance between adjacent data points (2-way ANOVA with Tukey’s correction). (B) Optomotor response in wild-type flies responding to gratings that were stationary (0 Hz) or rotating at 0.5 Hz, 3 Hz, 7 Hz, or 16 Hz. Higher optomotor index (OI) indicates greater optomotor response. n = 10. Asterisks indicate significance between adjacent points (A), significance via 2-way ANOVA with Tukey’s correction. * p < .05; ** p< .01; *** p< .001. Error bars show S.E.M.

**Figure S2.**
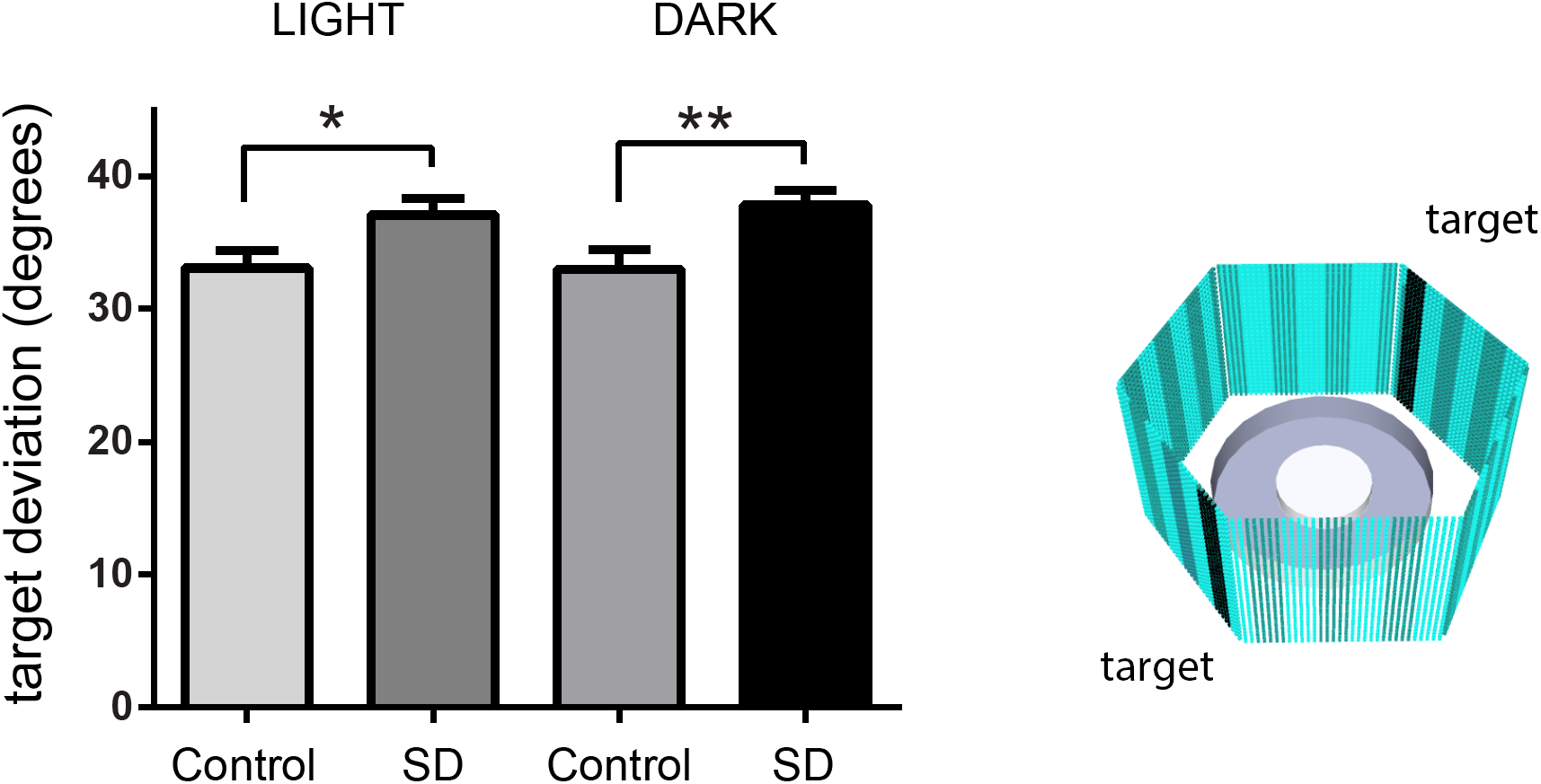
Sleep deprivation effects on attention are independent of visual experience. Target deviation during figure/ground discrimination in flies that were sleep deprived for 24 hrs in normal circadian conditions (light) or in 24 hrs of darkness (dark). The visual stimuli were a 7 Hz flickering object (’target’) and a mid luminance contrast grating (see methods). n>19 flies per data set. *p=0.03,**p=0.01 by t-test. Error bars indicate the s.e.m.

